# Differential Disease Severity and Whole Genome Sequence Analysis for Human Influenza A/H1N1pdm Virus in 2015-2016 Influenza Season

**DOI:** 10.1101/2020.02.20.957068

**Authors:** Hsuan Liu, Yu-Nong Gong, Kathryn Shaw-Saliba, Thomas Mehoke, Jared Evans, Zhen-Ying Liu, Mitra Lewis, Lauren Sauer, Peter Thielen, Richard Rothman, Kuan-Fu Chen, Andrew Pekosz

## Abstract

During the 2015-16 winter, the US experienced a relatively mild influenza season compared to Taiwan which had a higher number of total and severe cases. While H1N1pdm viruses dominated global surveillance efforts that season, the global distribution of genetic variants and their contributions to disease severity have not been investigated. Samples collected from influenza A positive patients by the Johns Hopkins Center of Excellence for Influenza Research and Surveillance (JH-CEIRS) active surveillance in the emergency rooms in Baltimore, Maryland, USA and northern Taiwan between November 2015 and April 2016, were processed for influenza A virus whole genome sequencing. In Baltimore, the majority of the viruses were the H1N1pdm clade 6B.1 and no H1N1pdm clade 6B.2 viruses were detected. In northern Taiwan, more than half of the H1N1pdm viruses were clade 6B.1 and 38% were clade 6B.2, consistent with the global observation that most 6B.2 viruses circulated in Asia and not North America. Whole virus genome sequence analysis identified two genetic subgroups present in each of the 6B.1 and 6B.2 clades and one 6B.1 intraclade reassortant virus. Clinical data showed 6B.2 patients had more disease symptoms including higher crude and inverse probability weighted odds of pneumonia than 6B.1 patients, suggesting 6B.2 circulation may contribute to the severe flu season in Taiwan. Local surveillance efforts linking H1N1pdm virus sequences to patient clinical and demographic data improve our understanding of influenza circulation and disease potential.

## Introduction

The 2009 pandemic H1N1 Influenza A virus (H1N1pdm) has supplanted the previous human H1N1 viruses to become the seasonal human H1N1 virus. In the 2015-16 influenza season, the northern hemisphere experienced its second global H1N1pdm-dominant year since 2009. The 2015-16 Influenza activity in the US was considered moderate, with lower numbers of total cases and a later peak epidemic when compared with the previous three influenza seasons [1]. However, in Taiwan, the 2015-16 season was much more serious [2], with an earlier start and higher numbers of total as well as severe, influenza cases compared to the prior influenza season 2014-15 [3].

Influenza A viruses (IAV) are subdivided into subtypes based on the antibody response to the viral glycoproteins, hemagglutinin (HA) and neuraminidase (NA). Viruses are further divided into genetic clades based on HA sequences in order to monitor for mutations that might lead to antigenic drift. The 2015-16 northern hemisphere vaccine strain of H1N1pdm, A/California/7/2009, is defined as H1 clade 1. Prior to the 2015-16 season, H1N1pdm was dominated by clade 6B. The global rise of H1 clade 6B.1 carrying amino acid changes S84N, S162N and I216T from clade 6B was detected in human surveillance efforts starting in June 2015 and is now the dominant H1N1 clade. The H1 clade 6B.2 viruses (V152T, V173I of HA1 and E164G and D174E of HA2) emerged in July 2015, peaked in January 2016, were found primarily in Asia and disappeared from surveillance efforts at the end of 2016 [4, 5]. The majority of 6B.1 and 6B.2 viruses were antigenically similar to the recommended components of the 2015-16 Northern Hemisphere H1N1pdm influenza vaccine A/California/7/2009 [1, 3, 6, 7], although antigenic difference between egg-adapted and circulating virus strains may have contributed to increased numbers of cases [8].

The genome of IAV consist of 8 segments of negative-sense RNA. In addition to HA mutations, mutations in the other 7 viral gene segments also result in genetic variants which can alter virus fitness, subvert pre-existing immunity and/or reduce vaccine effectiveness. Any of these changes could potentially alter disease severity or total number of cases [9]. Next-generation sequencing provides a sensitive and rapid method for sequencing of viruses directly from human specimens without any need for virus isolation or culture – both of which are known to select for viruses bearing additional mutations [10].

Most current global influenza surveillance systems lack connections between viral sequences and detailed patient demographic and clinical data. The Johns Hopkins Center of Excellence for Influenza Research and Surveillance (JH-CEIRS) has been actively performing human influenza surveillance since 2014. Surveillance efforts involve the connection of viral genome data and isolates with patient demographic and clinical data to gain more insights into how virus genetic variation can affect clinical disease. We hypothesized that sequence differences in influenza viruses could be associated with distinct clinical outcomes. This study involved concurrent, active influenza surveillance during the 2015-16 season in Baltimore, USA and northern Taiwan to compare and contrast differential epidemics in the same year between the two geographic regions and to analyze possible disease severity with influenza viral genotypes.

## Materials and Methods

### Active surveillance enrollment and sample collection

Institutional Review Boards at the Johns Hopkins University School of Medicine and Chang Gung Memorial Hospital provided ethical approval of the study (IRB00052743). From November 2015 to April 2016, active surveillance was performed in the adult emergency departments of the Johns Hopkins Medical Institutions (JHMI) at the East Baltimore and Bayview campuses in Baltimore, Maryland, USA and Chang Gung Memorial Hospitals (CGMH) in the northern Taiwan metropolitan area, including hospitals in the Taipei, Linkou, and Keelung Branches. Influenza-like illness was defined as documented or reported ≥38°C fever and any of one of three respiratory symptoms including cough, headache and/or sore throat within the past 7 days. Exclusion criteria included subjects who are unable to speak and understand English (in US) or Mandarin (in Taiwan), unable to provide written informed consent, currently incarcerated, or previously enrolled in the study during the same influenza season. Patients were approached by trained clinical coordinators who obtained written, informed consent before collected specimens and demographic and clinical data using a standard questionnaire. Data was confirmed by examination of the patient’s electronic health record. All data was de-identified and stored in a secure REDCap database [11, 12].

### Diagnosis and subtyping

Nasopharyngeal swabs or nasal washes of patients collected at the time of enrollment were tested for influenza A virus infection using the Cepheid Xpert Flu/RSV test (Cepheid, Sunnyvale, CA). Samples that were influenza A virus positive were further subtyped by qRT-PCR with specific H1 or H3 primers and probes according to USA Centers for Disease Control and Prevention (CDC) protocols. Only H1N1pdm positive samples were moved on to whole genome sequencing (WGS) analysis.

### Whole genome sequencing

Viral RNA was isolated from the influenza A positive clinical samples using the MagMax Viral RNA isolation reagent (Thermo Fisher) on an Eppendorf epMotion 5073 liquid handling workstation. Samples were processed for WGS using multi-segment PCR [10] and then prepared for deep sequencing using the Nextera XT library preparation reagent (Illumina). Samples were sequenced on the Illumina NextSeq platform. Raw paired-end data were first processed through Trim galore! (https://www.bioinformatics.babraham.ac.uk/projects/trim_galore/) with a quality score of 30 and an adapter sequence of CTGTCTCTTATACACATCT, only retaining pairs that both passed through this quality control step. The quality-trimmed reads were then aligned to the influenza vaccine strain reference sequence using bowtie2 (version 2.1.0) [13] with the ‘--very-sensitive-local’ option and converted to sorted BAM files using samtools [14]. Only sequences of all 8 gene segments passed quality controls were included in the study. The sequences were submitted to NCBI GenBank database and accession numbers are listed in the Table S1.

### Phylogenetics and sequence analysis

73 H1N1pdm clinical samples which yielded sufficient WGS coverage of the IAV genome were included in sequence analysis. In addition to these genomes, we retrieved two reference genomes of vaccine strains from the Global Initiative on Sharing All Influenza Data (GISAID, https://www.gisaid.org/) for further analyses. Nucleotide sequences of the longest open reading frames (ORFs) in each gene segment were used to build phylogenetic trees, including PB2, PB1, PA, HA, NP, NA, M1, and NS1, as well as, their alternative splicing (M2 and NS2) and ribosomal frameshift (PA-X) products. Nucleotide sequences of these ORFs (except for PA-X) in each strain were further concatenated to generate one single sequence for constructing a whole-genome phylogenetic tree. Phylogenetic trees for investigating their genetic or genomic relationships were based on maximum likelihood (ML) analysis, generated with RAxML (version 8.2.12) [15] under the GTRGAMMA model with 1000 bootstrap replicates. Their time scales based on sample collection dates were calibrated using the coalescent model in TreeTime [16] after building ML trees. Clade 6B, 6B.1, and 6B.2 were defined by the rules of HA amino acid mutations of global surveillance groups from nextflu [4]. Phylogenetic groups in other gene segments were based on their tree branches to separate annotated clades (e.g., 6B.1 and 6B.2). Inconsistent positioning of strains, or clade groups on individual trees was used to identify influenza reassortment. Individual gene segment clades were annotated and visualized using the ggtree R package [17].

All H1N1pdm genomes (n=2873) isolated from September, 2015 to May, 2016 were downloaded from GISAID as of January 2020, and were further aligned using MAFFT tool [18] under the default conditions. Moreover, we retrieved all NP sequences (n=20,477) of H1N1pdm isolated after the year of 2009 (inclusive) from GISAID as of January 2020 to investigate the population of viruses encoding an upstream translation initiation codon (AUG) in the NP gene segment. 20,470 NP sequences were used for further analyses, after aligning and removing 7 sequences with unexpected gaps.

### Glycosylation prediction

Potential N-linked glycosylation sites of the 6B.1 and 6B.2 HA and NA proteins were predicted by the NetNGlyc 1.0 server (http://www.cbs.dtu.dk/services/NetNGlyc).

### Statistical analysis of demographic and clinical data

To identify a contribution of virus clades and genetic subgroups to clinical symptoms, we applied Chi-square or Fisher’s exact tests for categorical variables and the rank-sum test for continuous variables when appropriate. The two-tailed statistical significance was set at p-value <0.05. When clinical or demographic category reached statistical significance in a univariate analysis, confounding effects were then analyzed. In addition to adjusting for possible confounding effects between demographics, disease severity and clinical outcomes, a propensity score-based weighting was applied. Briefly, we applied those potential confounders to generate the propensity (or probability) of being infected with different subgroups of H1N1pdm viruses by using machine learning based modeling methods [19]. By using the inverse probability of treatment weighting (IPTW) method, patients with higher propensity will have lower weights in the final logistic regression model to adjust for confounding factors. The weights create a pseudo-population where the weighted virus subgroups have similar distribution across these confounders. In one simulation model, the propensity score-based weighting methods could adjust for most of the confounding effects [20]. Odds ratios of unadjusted and IPTW were calculated using logistic regression analysis.

## Results

### H1N1pdm clade 6B.1 predominantly circulated in Baltimore and clade 6B.1 and 6B.2 co-circulated in northern Taiwan

From November 2015 to April 2016, 261 and 256 symptomatic patients were enrolled in the study from emergency departments in JHMI (Baltimore) and CGMH (northern Taiwan). IAV was found in 32.6% (85/261) of symptomatic patients from Baltimore and 57.8% (148/256) from northern Taiwan. About half of the IAV-positive samples were further subtyped, and the demographics and comorbidities were found to be similar. In 97.5% (39/40) of samples from Baltimore and 90.1% (64/71) of samples from northern Taiwan H1N1 was the influenza A virus subtype. The epidemic curves of the two surveillance sites were different, with influenza A virus activity starting in January and peaking in mid-March in Baltimore while IAV activity in northern Taiwan started in November and peaked in late January/early February, with peak activity higher than in Baltimore (Fink et al., MedRxiv).

Seventy-three of the 103 H1N1pdm clinical samples yielded sufficient sequence coverage of the IAV genome. Virus samples were initially analyzed based on HA segment sequences (Figure 1). In Baltimore, 91.7% (22/24) of viruses were clade 6B.1, two were clade 6B, and no 6B.2 viruses were identified. The northern Taiwan samples consisted of 57.1% (28/49) clade 6B.1, 38.8% (19/49) clade 6B.2 and 4.1% (2/49) clade 6B viruses. The difference in clade circulation is consistent with global influenza surveillance that H1N1pdm 6B.1 and 6B.2 were dominant in the 2015-16 season and most 6B.2 viruses were found in Asia [5, 21].

**Figure 1.**
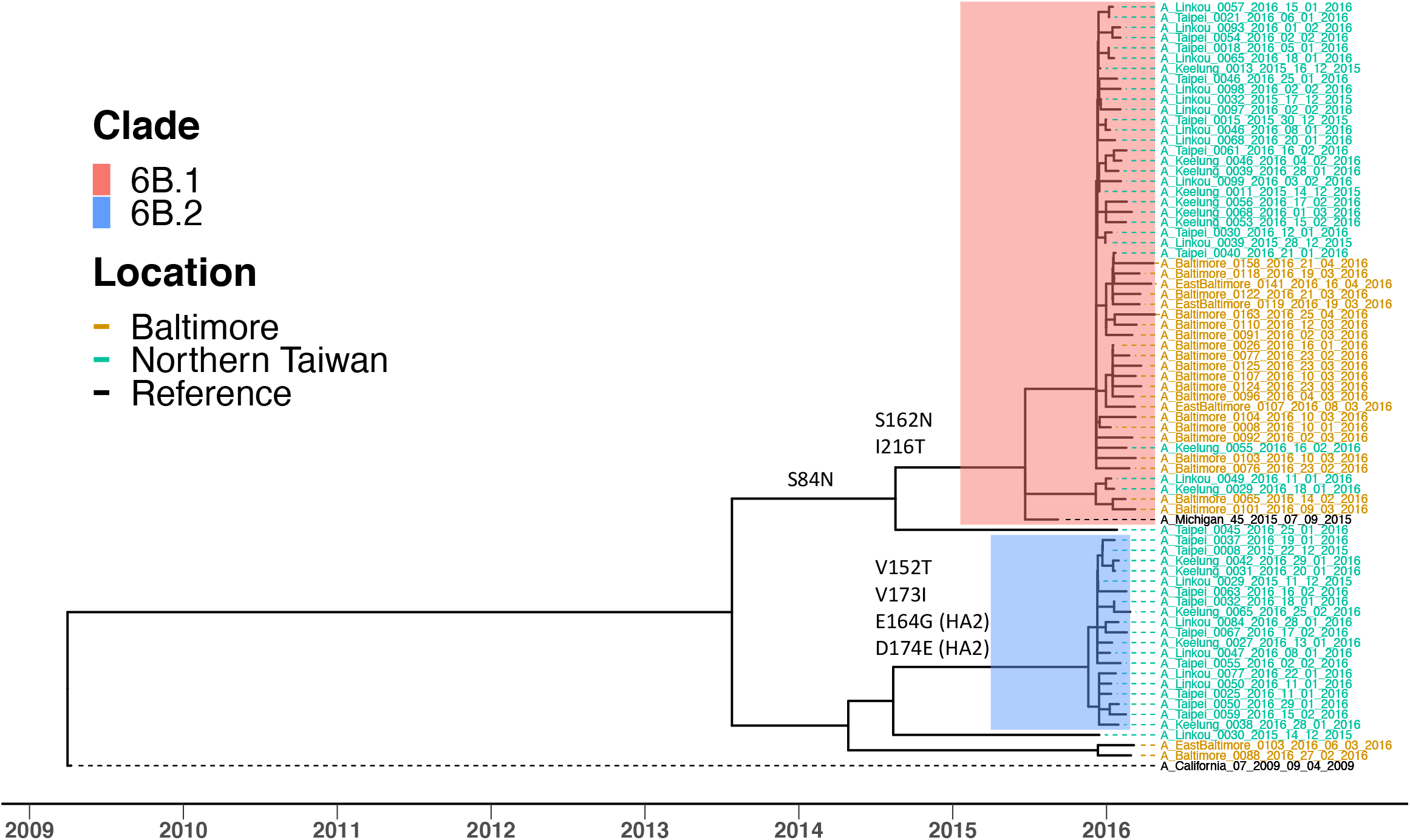
H1N1pdm clade 6B.1 and clade 6B.2 circulated in Baltimore and northern Taiwan in the 2015-16 season. Maximum likelihood (ML) tree of HA gene coding sequences from 73 H1N1pdm viruses of the surveillance was generated using RAxML under GTRGAMMA model with 1000 bootstrap replicates. Their time scales based on sample collection dates, noted in each sample, were calibrated using coalescent model in TreeTime after building ML tree. Reference sequences are H1N1pdm vaccine strains, A/California/07/2009 and A/Michigan/45/2015. Tips were colored by location of sample collection. Samples from Baltimore were colored in brown; samples from northern Taiwan were colored in green. Most samples were grouped into two clusters, 6B.1 (in red) and 6B.2 (in blue). Specific amino acid mutations defining major branches of 6B.1 and 6B.2 were indicated on the side of branch.

### Identification of four distinct genetic subgroups of H1N1pdm

To better understand the genetic diversity in the identified HA clades, we performed a phylogenetic analysis using concatenated gene segments (Figure 2) and individual ORF trees (Figure S1 and Figure 2). Branch tips of the 10 ORFs from the other seven gene segments were colored by the HA clade of each sample and defining amino acid mutations were noted (Figure S1). For the most part, the sequences of other ORFs clustered together consistent with their HA clade (Figure 2). Amino acid differences between clades 6B.1 and 6B.2 are listed for HA (Table S2), NA (Table S3) and the other ORFs (Table S4).

**Figure 2.**
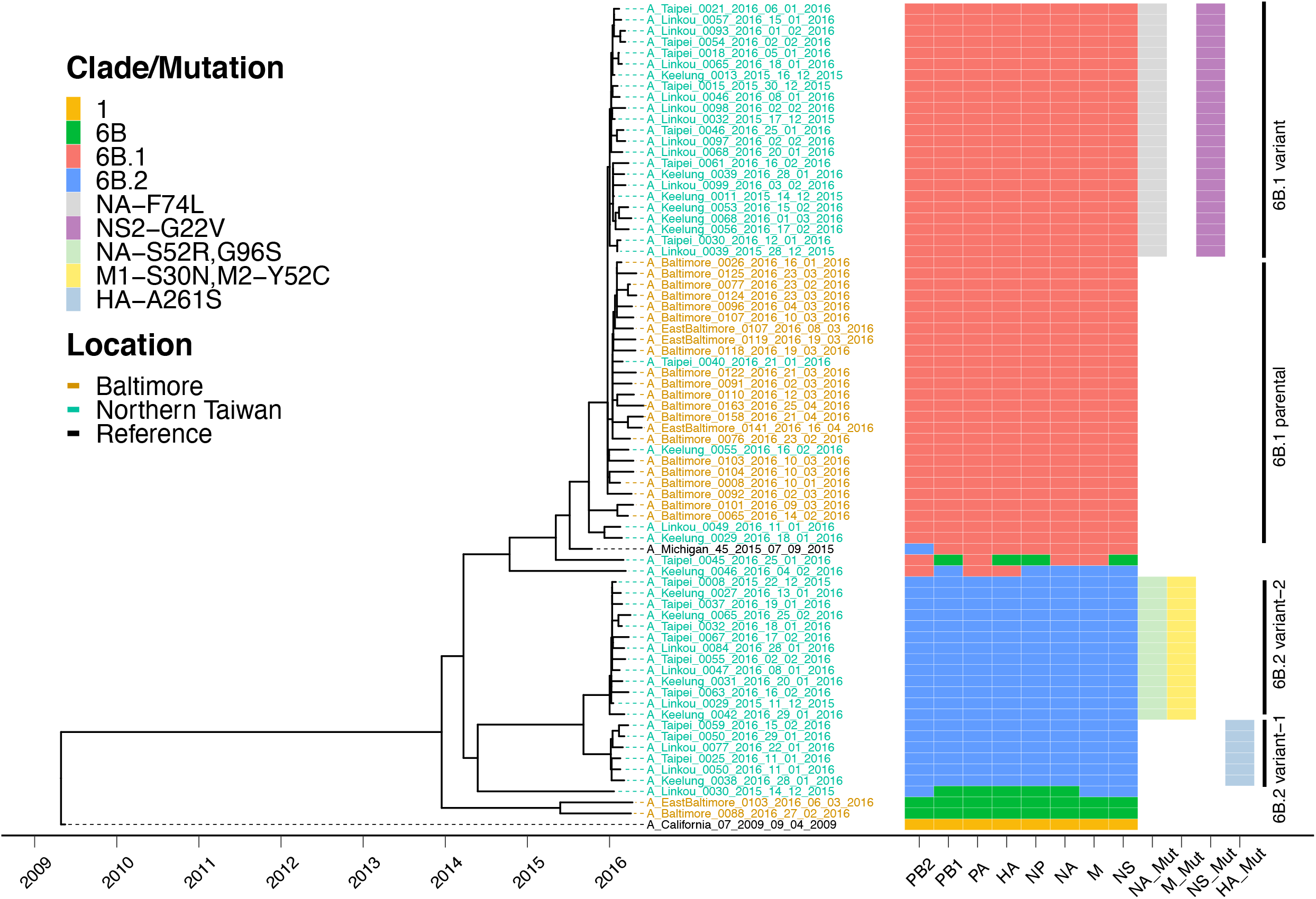
Four distinct genetic subgroups of H1N1pdm were identified in Baltimore and northern Taiwan in the 2015-16 season. Maximum likelihood (ML) trees of concatenated gene segments of 73 H1N1pdm viruses of the surveillance were generated using RAxML under GTRGAMMA model with 1000 bootstrap replicates. Their time scales based on sample collection dates, noted in each sample, were calibrated using coalescent model in TreeTime after building ML trees. Reference sequences are H1N1pdm vaccine strains, A/California/07/2009 and A/Michigan/45/2015. Tips were colored by location of sample collection. Samples from Baltimore were colored in brown; samples from northern Taiwan were colored in green. A schematic representation of clade/mutation of each gene segment based on their individual gene tree branches to separate annotated clades (1,6B, 6B.1, or 6B.2) and amino acid mutations (see Figure S1) was annotated and visualized using the ggtree R package. Mutations (Mut) in NA, M, NS, and HA segments were identified to be able to divide majority of viruses into four distinct genetic subgroups, 6B.1 parental, 6B.1 variant, 6B.2 variant-1 and 6B.2 variant-2.

Additional amino acid mutations were identified to differentiate viruses into four distinct virus genetic subgroups (Figure 2). The 6B.1 viruses were separated into two groups, a 6B.1 parental and a 6B.1 variant defined by unique amino acid changes in NA and NS2 (Table S3 and S4). All Baltimore 6B.1 viruses were 6B.1 parental. In northern Taiwan, only 4 were 6B.1 parental, with the remaining clade 6B.1 viruses (23/27) being 6B.1 variant. The 6B.2 viruses were also separated into two genetic subgroups - variant-1 and variant-2 – that are differentiated by mutations in HA, NA, M1 and M2 (Table S2, S3 and S4).

### Dynamic virus evolution of gene segments and possible reassortment

All segments in two of the four 6B viruses, A/EastBaltimore/0103/2016 and A/Baltimore/0088/2016, were clade 6B. One 6B virus A/Taipei/0045/2016 contained mutations in PB2, PA, NA and M segments consistent with those found in the clade 6B.1 and the remaining segments were similar but distinct from clade 6B.1. The other 6B virus A/Linkou/0030/2015 contained sequences in the PB2, M and NS segments consistent with the clade 6B.2 viruses and the remaining segments were related to but distinct from clade 6B.2. (Figure 2 and S1). These suggested that all segments of 6B viruses were evolving toward either the clade 6B.1 or clade 6B.2.

There was also evidence of intraclade reassortment in one northern Taiwan 6B.1 sample A/Keelung/0046/2016, as three gene segments of 6B.1 (segments HA, PB2 and PA) and five 6B.2 (segments PB1, NP, NA, M and NS) were detected (Figure 2). While we cannot rule out co-infection with two H1N1 clade viruses, given the number of reads present for each segment, it seems more likely this individual was infected with an intraclade reassortant virus.

### HA and NA N-linked glycosylation

Eight N-linked glycosylation sites on the HA of A/California/7/2009 were previously determined by sequence analysis and verified by mass spectrometry [22]. The 6B.1 HA sequence predicts an additional N-linked glycosylation site at 162-164 (N-Q-S) on the Sa antigenic site of the head domain due to a S to N mutation at HA-162 (Table S2) [23]. The NA from the A/California/7/2009 vaccine strain has 8 potential N-linked glycosylation sites and 6 sites were experimentally confirmed [22]. Both 6B.1 and 6B.2 viruses gain one additional predicted N-linked glycosylation site at 42-44 (N-Q-S) of the NA stalk and lose one at 386-388 (N-F-S) which lies on the lateral surface of the NA head. In addition, 68.4% (13/19) of 6B.2 viruses with NA-S52R mutation lose one glycosylation site at 50-52 (N-Q-S) on the NA stalk (Table S3).

### Antiviral resistance mutations

All sequences contained the amantadine resistance mutation M2-S31N which is found in the A/Cal/07/2009 virus. One 6B.2 variant-2 sample from northern Taiwan, A/Linkou/0029/2015, contained the NA-H275Y, the most common mutation associated with oseltamivir resistance in N1 NA.

### An upstream AUG codon lengthens the nucleoprotein (NP) of H1N1pdm

The NP gene segments of all samples in the study have a new, in frame AUG start codon 18 nucleotides upstream of the usual start codon of NP proteins of IAV including human seasonal H1N1 and H3N2 [24, 25], adding 6 amino acids to the amino terminus of the protein with clade 6B.1 (M-S-D-I-E-A) and 6B.2 viruses (M-S-D-I-E-V) differing at amino acid six. This start codon was shown to be utilized and the resulting NP protein capable of supporting viral polymerase activity [26, 27]. The presence of an upstream NP start codon and the amino acid difference at position 6 between 6B.1 and 6B.2 were also present in approximately 70% of GISAID sequences (n=20,470) of H1N1pdm NP since 2009 with an Alanine (A) dominating at position 6.

### Patients infected with clade 6B.2 had increased symptoms of severe influenza

To determine whether viral genotype contributes to symptoms and disease severity, the demographic and clinical data from patients infected with clade 6B.1 or 6B.2 viruses were first compared by univariate analyses (Table 1). Patients infected with 6B.2 were more likely to be older, female, Asian, and unvaccinated. These patients had a higher prevalence of runny nose and diarrhea and had increased symptoms of severe influenza, including increased likelihood of oxygen supplementation, longer in-hospital stay, and pneumonia diagnosis (by radiological findings). Interestingly, patients infected with clade 6B.1 viruses had a lower number of co-morbidities associated with severe influenza, suggesting a healthier population was being infected with those viruses.

**Table 1.**
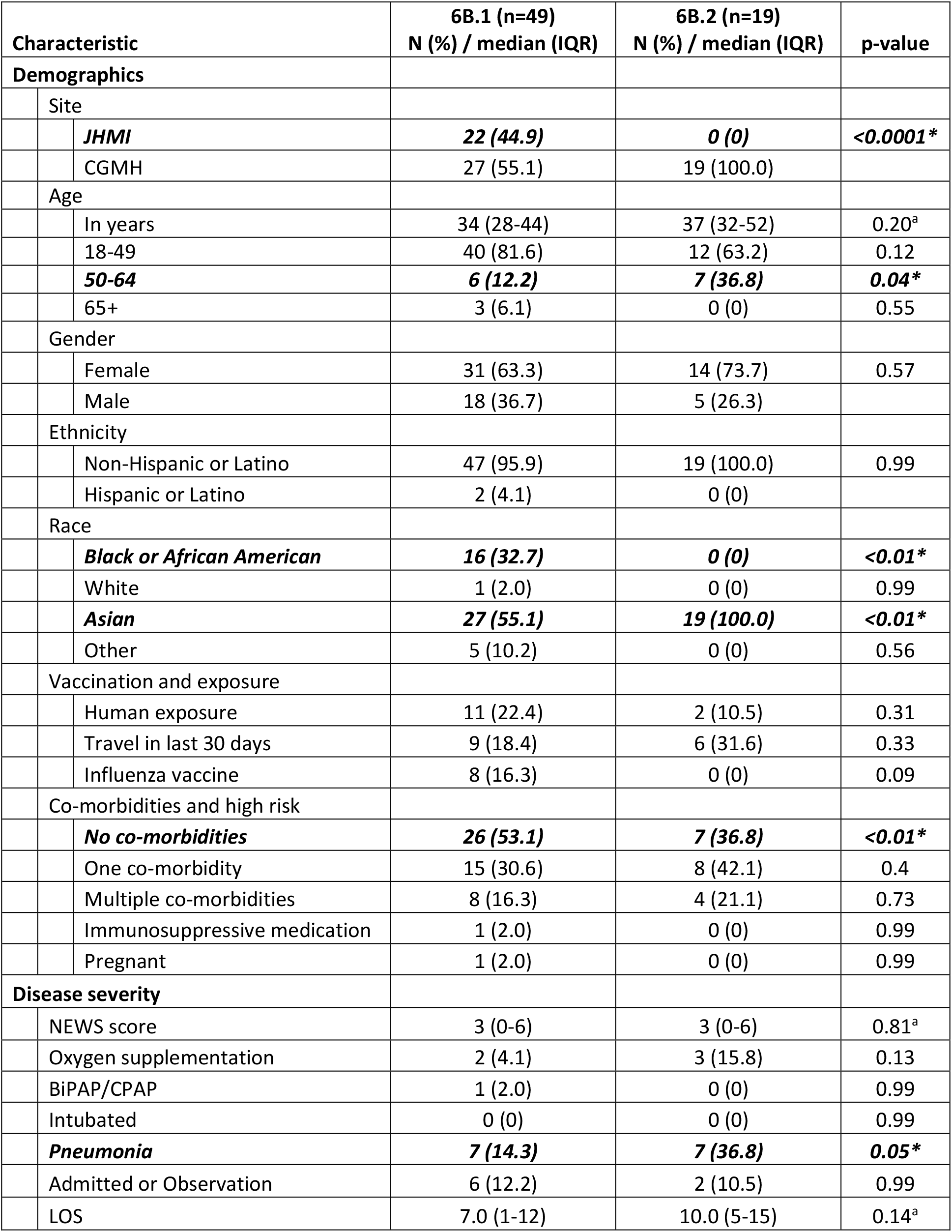
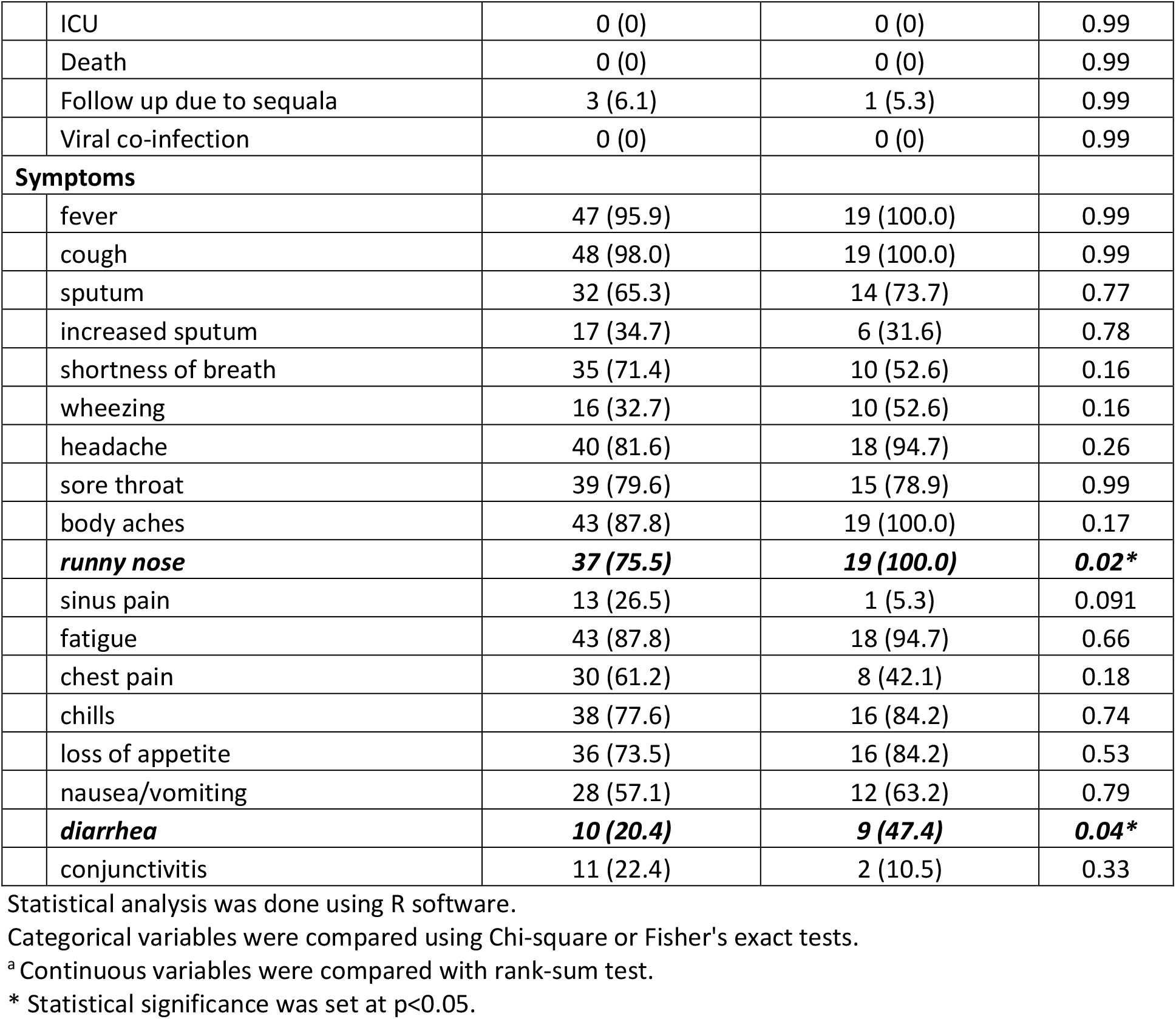
Univariate analysis of patients infected with H1N1pdm 6B.1 and 6B.2 viruses of Baltimore and northern Taiwan in the 2015-16 season

| Characteristic |  | 6B.1 (n=49)<br>N (%) / median (IQR) | 6B.2 (n=19)<br>N (%) / median (IQR) | p-value |
| --- | --- | --- | --- | --- |
| **Demographics** |  |  |  |  |
|  | Site |  |  |  |
|  | *JHMI* | **22 (44.9)** | **0 (0)** | **<0.0001\*** |
|  | CGMH | 27 (55.1) | 19 (100.0) |  |
|  | Age |  |  |  |
|  | In years | 34 (28-44) | 37 (32-52) | 0.20<sup>a</sup> |
|  | 18-49 | 40 (81.6) | 12 (63.2) | 0.12 |
|  | **50-64** | **6 (12.2)** | **7 (36.8)** | **0.04\*** |
|  | 65+ | 3 (6.1) | 0 (0) | 0.55 |
|  | Gender |  |  |  |
|  | Female | 31 (63.3) | 14 (73.7) | 0.57 |
|  | Male | 18 (36.7) | 5 (26.3) |  |
|  | Ethnicity |  |  |  |
|  | Non-Hispanic or Latino | 47 (95.9) | 19 (100.0) | 0.99 |
|  | Hispanic or Latino | 2 (4.1) | 0 (0) |  |
|  | Race |  |  |  |
|  | **Black or African American** | **16 (32.7)** | **0 (0)** | **<0.01\*** |
|  | White | 1 (2.0) | 0 (0) | 0.99 |
|  | **Asian** | **27 (55.1)** | **19 (100.0)** | **<0.01\*** |
|  | Other | 5 (10.2) | 0 (0) | 0.56 |
|  | Vaccination and exposure |  |  |  |
|  | Human exposure | 11 (22.4) | 2 (10.5) | 0.31 |
|  | Travel in last 30 days | 9 (18.4) | 6 (31.6) | 0.33 |
|  | Influenza vaccine | 8 (16.3) | 0 (0) | 0.09 |
|  | Co-morbidities and high risk |  |  |  |
|  | **No co-morbidities** | **26 (53.1)** | **7 (36.8)** | **<0.01\*** |
|  | One co-morbidity | 15 (30.6) | 8 (42.1) | 0.4 |
|  | Multiple co-morbidities | 8 (16.3) | 4 (21.1) | 0.73 |
|  | Immunosuppressive medication | 1 (2.0) | 0 (0) | 0.99 |
|  | Pregnant | 1 (2.0) | 0 (0) | 0.99 |
| **Disease severity** |  |  |  |  |
|  | NEWS score | 3 (0-6) | 3 (0-6) | 0.81<sup>a</sup> |
|  | Oxygen supplementation | 2 (4.1) | 3 (15.8) | 0.13 |
|  | BiPAP/CPAP | 1 (2.0) | 0 (0) | 0.99 |
|  | Intubated | 0 (0) | 0 (0) | 0.99 |
|  | **Pneumonia** | **7 (14.3)** | **7 (36.8)** | **0.05\*** |
|  | Admitted or Observation | 6 (12.2) | 2 (10.5) | 0.99 |
|  | LOS | 7.0 (1-12) | 10.0 (5-15) | 0.14<sup>a</sup> |
|  | ICU | 0 (0) | 0 (0) | 0.99 |
|  | Death | 0 (0) | 0 (0) | 0.99 |
|  | Follow up due to sequela | 3 (6.1) | 1 (5.3) | 0.99 |
|  | Viral co-infection | 0 (0) | 0 (0) | 0.99 |
| **Symptoms** |  |  |  |  |
|  | fever | 47 (95.9) | 19 (100.0) | 0.99 |
|  | cough | 48 (98.0) | 19 (100.0) | 0.99 |
|  | sputum | 32 (65.3) | 14 (73.7) | 0.77 |
|  | increased sputum | 17 (34.7) | 6 (31.6) | 0.78 |
|  | shortness of breath | 35 (71.4) | 10 (52.6) | 0.16 |
|  | wheezing | 16 (32.7) | 10 (52.6) | 0.16 |
|  | headache | 40 (81.6) | 18 (94.7) | 0.26 |
|  | sore throat | 39 (79.6) | 15 (78.9) | 0.99 |
|  | body aches | 43 (87.8) | 19 (100.0) | 0.17 |
|  | **runny nose** | **37 (75.5)** | **19 (100.0)** | **0.02\*** |
|  | sinus pain | 13 (26.5) | 1 (5.3) | 0.091 |
|  | fatigue | 43 (87.8) | 18 (94.7) | 0.66 |
|  | chest pain | 30 (61.2) | 8 (42.1) | 0.18 |
|  | chills | 38 (77.6) | 16 (84.2) | 0.74 |
|  | loss of appetite | 36 (73.5) | 16 (84.2) | 0.53 |
|  | nausea/vomiting | 28 (57.1) | 12 (63.2) | 0.79 |
|  | **diarrhea** | **10 (20.4)** | **9 (47.4)** | **0.04\*** |
|  | conjunctivitis | 11 (22.4) | 2 (10.5) | 0.33 |

Statistical analysis was done using R software.
Categorical variables were compared using Chi-square or Fisher's exact tests.
<sup>a</sup> Continuous variables were compared with rank-sum test.
\* Statistical significance was set at p<0.05.

We further utilized a logistic regression analysis to estimate the propensity of being infected by 6B.2, using potential confounders such as age, gender, comorbidities, obesity, vaccine status, human exposure and travel history (area under the ROC curve: 0.798). After IPTW to adjust for the possible confounding effects, infection with clade 6B.2 viruses was found to be significantly associated with pneumonia (Odds ratio: 3.261, 95% CI: 1.696-4.826, p=0.008) (Table 2).

**Table 2.** Logistic regression analysis of pneumonia in patients infected with H1N1pdm 6B.1 and 6B.2 viruses of Baltimore and northern Taiwan in the 2015-16 season

|  |  | Unadjusted |  |  | IPTW** |  |  |
| --- | --- | --- | --- | --- | --- | --- | --- |
| Clade | Pneumonia | Odds Ratio | 95% CI | p-value | Odds Ratio | 95% CI | p-value |
| 6B.1 | 14.3% (7/49) | 1 | - | - | 1 | - | - |
| 6B.2 | 36.8% (7/19) | 3.500 | 1.629-5.372 | 0.046* | 3.261 | 1.696-4.826 | 0.008* |

\*p<0.05, logistic regression analysis.
\*\* IPTW (inverse probability of treatment weighting) results were done by propensity score weighting with recursive partitioning algorithm (area under the ROC curve: 0.798) for age, gender, comorbidities, obesity, vaccine status, human exposure and travel history, then followed by logistic regression analysis.

The distribution of cases across the four genetic subgroups was then determined (Tables 3 and 4). Patients infected with the 6B.1 parental virus had increased reporting of wheezing, sinus pain, and nausea/vomiting, along with the expected difference in race and ethnicity stemming from the identification of the 6B.1 variant only in northern Taiwan (Table 3). When patients infected with the 6B.2 variant-1 and variant-2 viruses were compared there were no obvious difference in symptoms (Table 4), however, the small sample sizes in the 6B.2 subgroups precluded a definitive characterization of demographic and clinical data between these genotypes. Overall, our data suggested clade 6B.2 circulation in northern Taiwan likely played a significant role in differentiating the influenza epidemics between Baltimore and northern Taiwan in the season.

**Table 3.** Univariate analysis of patients infected with H1N1pdm 6B.1 parental and 6B.1 variant viruses of Baltimore and northern Taiwan in the 2015-16 season

| Characteristic |  | 6B.1 Parental (n=26)<br>N (%) / median (IQR) | 6B.1 Variant (n=23)<br>N (%) / median (IQR) | p-value |
| --- | --- | --- | --- | --- |
| **Demographics** |  |  |  |  |
|  | Site |  |  |  |
|  | *JHMI* | **22 (84.6)** | **0 (0)** | **<0.0001\*** |
|  | CGMH | 4 (15.4) | 23 (100.0) |  |
|  | Age |  |  |  |
|  | In years | 33 (29-44) | 35 (29-44) | 0.75<sup>a</sup> |
|  | 18-49 | 20 (76.9) | 20 (87.0) | 0.47 |
|  | 50-64 | 5 (19.2) | 1 (4.3) | 0.19 |
|  | 65+ | 1 (3.8) | 2 (8.7) | 0.59 |
|  | Gender |  |  |  |
|  | Female | 14 (53.8) | 17 (73.9) | 0.24 |
|  | Male | 12 (46.2) | 6 (26.1) |  |
|  | Ethnicity |  |  |  |
|  | Non-Hispanic or Latino | 24 (92.3) | 23 (100.0) | 0.49 |
|  | Hispanic or Latino | 2 (7.7) | 0 (0) |  |
|  | Race |  |  |  |
|  | **Black or African American** | **16 (61.5)** | **0 (0)** | **<0.0001\*** |
|  | White | 1 (3.8) | 0 (0) | 0.99 |
|  | **Asian** | **4 (15.4)** | **23 (100.0)** | **<0.0001\*** |
|  | Other | 5 (19.3) | 0 (0) | 0.08 |
|  | Vaccination and exposure |  |  |  |
|  | Human exposure | 4 (15.4) | 7 (30.4) | 0.36 |
|  | Travel in last 30 days | 3 (11.5) | 6 (26.1) | 0.35 |
|  | Influenza vaccine | 6 (23.1) | 2 (8.7) | 0.33 |
|  | Co-morbidities and high risk |  |  |  |
|  | No co-morbidities | 11 (42.3) | 15 (65.2) | 0.15 |
|  | One co-morbidity | 8 (30.8) | 7 (30.4) | 0.99 |
|  | Multiple co-morbidities | 7 (26.9) | 1 (4.3) | 0.05 |
|  | Immunosuppressive medication | 1 (3.8) | 0 (0) | 0.99 |
|  | Pregnant | 1 (3.8) | 0 (0) | 0.99 |
| **Disease severity** |  |  |  |  |
|  | NEWS score | 3 (0-6) | 3 (0-6) | 0.57<sup>a</sup> |
|  | Oxygen supplementation | 2 (7.7) | 0 (0) | 0.49 |
|  | BiPAP/CPAP | 1 (3.8) | 0 (0) | 0.49 |
|  | Intubated | 0 (0) | 0 (0) | 0.99 |
|  | Pneumonia | 3 (11.5) | 4 (17.4) | 0.69 |
|  | Admitted or Observation | 4 (15.4) | 2 (8.7) | 0.67 |
|  | LOS | 6.5 (1-12) | 6.0 (3-9) | 0.67<sup>a</sup> |
|  | ICU | 0 (0) | 0 (0) | 0.99 |
|  | Death | 0 (0) | 0 (0) | 0.99 |
|  | Follow up due to sequela | 2 (7.7) | 1 (4.3) | 0.99 |
|  | Viral co-infection | 0 (0) | 0 (0) | 0.99 |
| **Symptoms** |  |  |  |  |
|  | fever | 24 (92.3) | 23 (100.0) | 0.49 |
|  | cough | 25 (96.2) | 23 (100.0) | 0.99 |
|  | sputum | 15 (57.7) | 17 (73.9) | 0.37 |
|  | increased sputum | 10 (38.5) | 7 (30.4) | 0.77 |
|  | shortness of breath | 20 (76.9) | 15 (65.2) | 0.53 |
|  | **wheezing** | **12 (46.2)** | **4 (17.4)** | **0.04\*** |
|  | headache | 23 (88.5) | 17 (73.9) | 0.27 |
|  | sore throat | 19 (73.1) | 20 (87.0) | 0.3 |
|  | body aches | 24 (92.3) | 19 (82.6) | 0.4 |
|  | runny nose | 17 (65.4) | 20 (87.0) | 0.1 |
|  | **sinus pain** | **11 (42.3)** | **2 (8.7)** | **0.01\*** |
|  | fatigue | 22 (84.6) | 21 (91.3) | 0.67 |
|  | chest pain | 19 (73.1) | 11 (47.8) | 0.08 |
|  | chills | 21 (80.8) | 17 (73.9) | 0.73 |
|  | loss of appetite | 20 (76.9) | 16 (69.6) | 0.75 |
|  | **nausea/vomiting** | **19 (73.1)** | **9 (39.1)** | **0.02\*** |
|  | diarrhea | 7 (26.9) | 3 (13.0) | 0.3 |
|  | conjunctivitis | 8 (30.8) | 3 (13.0) | 0.18 |

Statistical analysis was done using R software.
Categorical variables were compared using Chi-square or Fisher's exact tests.
<sup>a</sup> Continuous variables were compared with rank-sum test.
\* Statistical significance was set at p<0.05.

**Table 4.** Univariate analysis of patients infected with H1N1pdm 6B.2 variant-1 and 6B.2 variant-2 viruses of Baltimore and northern Taiwan in the 2015-16 season

| Characteristic |  | 6B.2 Variant-1 (n=6)<br>N (%) / median (IQR) | 6B.2 Variant-2 (n=13)<br>N (%) / median (IQR) | p-value |
| --- | --- | --- | --- | --- |
| **Demographics** |  |  |  |  |
|  | Site |  |  |  |
|  | JHMI | 0 (0) | 0 (0) | 0.99 |
|  | CGMH | 6 (100.0) | 13 (100.0) |  |
|  | Age |  |  |  |
|  | In years | 34 (29-48) | 44 (34-54) | 0.20<sup>a</sup> |
|  | 18-49 | 4 (66.7) | 8 (61.5) | 0.99 |
|  | 50-64 | 2 (33.3) | 5 (38.5) | 0.99 |
|  | 65+ | 0 (0) | 0 (0) | 0.99 |
|  | Gender |  |  |  |
|  | Female | 3 (50.0) | 11 (84.6) | 0.26 |
|  | Male | 3 (50.0) | 2 (15.4) |  |
|  | Ethnicity |  |  |  |
|  | Non-Hispanic or Latino | 6 (100.0) | 13 (100.0) | 0.99 |
|  | Hispanic or Latino | 0 (0) | 0 (0) |  |
|  | Race |  |  |  |
|  | Black or African American | 0 (0) | 0 (0) | 0.99 |
|  | White | 0 (0) | 0 (0) | 0.99 |
|  | Asian | 6 (100.0) | 13 (100.0) | 0.99 |
|  | Other | 0 (0) | 0 (0) | 0.99 |
|  | Vaccination and exposure |  |  |  |
|  | Human exposure | 0 (0) | 2 (15.4) | 0.83 |
|  | Travel in last 30 days | 3 (50.0) | 3 (23.1) | 0.32 |
|  | Influenza vaccine | 0 (0) | 0 (0) | 0.99 |
|  | Co-morbidities and high risk |  |  |  |
|  | No co-morbidities | 3 (50.0) | 4 (30.8) | 0.77 |
|  | One co-morbidity | 2 (33.3) | 6 (46.2) | 0.99 |
|  | Multiple co-morbidities | 1 (16.7) | 3 (23.1) | 0.99 |
|  | Immunosuppressive medication | 0 (0) | 0 (0) | 0.99 |
|  | Pregnant | 0 (0) | 0 (0) | 0.99 |
| **Disease severity** |  |  |  |  |
|  | NEWS score | 4 (2-6) | 3 (0-5) | 0.54<sup>a</sup> |
|  | Oxygen supplementation | 0 (0) | 3 (23.1) | 0.52 |
|  | BiPAP/CPAP | 0 (0) | 0 (0) | 0.99 |
|  | Intubated | 0 (0) | 0 (0) | 0.99 |
|  | Pneumonia | 2 (33.3) | 5 (38.5) | 0.99 |
|  | Admitted or Observation | 0 (0) | 2 (15.4) | 0.99 |
|  | **LOS** | **0 (0-0)** | **10.0 (5-15)** | **<0.01\*** |
|  | ICU | 0 (0) | 0 (0) | 0.99 |
|  | Death | 0 (0) | 0 (0) | 0.99 |
|  | Follow up due to sequela | 0 (0) | 1 (7.7) | 0.99 |
|  | Viral co-infection | 0 (0) | 0 (0) | 0.99 |
| **Symptoms** |  |  |  |  |
|  | fever | 6 (100.0) | 13 (100.0) | 0.99 |
|  | cough | 6 (100.0) | 13 (100.0) | 0.99 |
|  | sputum | 5 (83.3) | 9 (69.2) | 0.99 |
|  | increased sputum | 3 (50.0) | 3 (23.1) | 0.32 |
|  | shortness of breath | 2 (33.3) | 8 (61.5) | 0.35 |
|  | wheezing | 2 (33.3) | 8 (61.5) | 0.35 |
|  | headache | 6 (100.0) | 12 (92.3) | 0.99 |
|  | sore throat | 6 (100.0) | 9 (69.2) | 0.26 |
|  | body aches | 6 (100.0) | 13 (100.0) | 0.99 |
|  | runny nose | 6 (100.0) | 13 (100.0) | 0.99 |
|  | sinus pain | 0 (0) | 1 (7.7) | 0.99 |
|  | fatigue | 6 (100.0) | 12 (92.3) | 0.99 |
|  | chest pain | 2 (33.3) | 6 (46.2) | 0.99 |
|  | chills | 5 (83.3) | 11 (84.6) | 0.99 |
|  | loss of appetite | 6 (100.0) | 10 (76.9) | 0.52 |
|  | nausea/vomiting | 3 (50.0) | 9 (69.2) | 0.62 |
|  | diarrhea | 4 (66.7) | 5 (38.5) | 0.35 |
|  | conjunctivitis | 0 (0) | 2 (15.4) | 0.99 |

Statistical analysis was done using R software.
Categorical variables were compared using Chi-square or Fisher's exact tests.
<sup>a</sup> Continuous variables were compared with rank-sum test.
\* Statistical significance was set at $p < 0.05$ .

## Discussion

Four distinct genetic subgroups of H1N1pdm viruses were circulating in Baltimore and northern Taiwan during the 2015-16 flu season. While the 6B.1 clade was circulating at both sites, some 6B.1 viruses in northern Taiwan had additional mutations which defined a genetic subgroup we called 6B.1 variant. The two 6B.1 subgroups of viruses did not differ drastically with respect to disease severity. The 6B.2 clade viruses were detected in northern Taiwan but not in Baltimore and consisted of two distinct genetic subgroups. Patients infected with 6B.2 virus showed more and stronger disease symptoms by both uni- and multivariate analysis. This difference in H1N1pdm clade circulation may have contributed to the more severe influenza season experienced in northern Taiwan compared to Baltimore in 2015-16. After adjusting and weighting for age, gender, comorbidities, obesity, vaccine status, human exposure and travel history, people infected with 6B.2 still had a higher odds ratio of pneumonia (Table 2). The data indicate that infection with clade 6B.2 viruses is associated with a higher disease severity at our surveillance sites independent of patient demographics and comorbidities. The presence of the 6B.2 clade in addition to low preexisting immunity to circulating H1N1 in northern Taiwan (reference Fink, 2020 medrxiv) may be the most likely reasons explaining the different numbers and severity of influenza cases between Baltimore and northern Taiwan. The only report for symptoms of 6B.1 compared to 6B.2 infection was seen in Israel, where infection with 6B.2 virus was associated with more vomiting and nausea compared to 6B.1-infected patients [28]. In our surveillance, we did see a higher percentage of 6B.2 than 6B.1 patients with vomiting and nausea, but this did not reach statistical significance (Table 1).

The H1N1pdm 6B.1 and 6B.2 viruses dominated the 2015-16 season in the Northern Hemisphere. However, numbers of cases and disease severity differed between geographic regions. The 2015-16 H1N1pdm season in the US and Canada were moderate compared to the prior season [1, 6]. On the other hand, influenza surveillance in Russia, UK, eastern Europe, the Middle East and Asia (including Taiwan) had increasing numbers of severe cases [3, 7, 29–33]. Reports also showed H1N1pdm impacted the southern hemisphere in the 2016 season, in particular, Brazil and Reunion Island [34–36].

Serological surveillance indicated that the circulating H1N1pdm strains in 2015-16 were antigenically similar to the vaccine A/California/7/2009 in the US, UK and many other countries [1, 6, 7, 32, 37, 38]. This suggests that the HA amino acid differences between the circulating 6B.1 or 6B.2 viruses and the vaccine strain were likely not able to explain differential epidemics in different regions of the year even though vaccination coverage might differ. Whether mutations in the other 7 viral genes contributed to the variable disease severity reported in 2015-16 has not been studied carefully. Some mutations in NA and M1 occurred at sites previously associated with altered virus transmission [39, 40], but our initial experiments on human nasal epithelial cell cultures did not detect any replication differences between clade 6B.1 and 6B.2 viruses (reference Fink, 2020 medrxiv).

We identified one 6B.2 patient in northern Taiwan had NA-H275Y mutation, which is associated with NA drug resistance. In another study of 2015-16 H1N1pdm in Taiwan, one clinical isolate with NA-H275Y was resistant to oseltamivir [41]. The data suggest sporadic identification of oseltamivir resistant H1N1pdm viruses in 2015-16 in Taiwan.

The clade 6B.1 parental genetic subgroup continued to circulate after 2015-16, and evolved into several 6B.1A subclades through August 2020. It is not clear what drove the extinction of 6B.2 but it is possible that clade 6B.1 viruses were better adapted to infect and spread in humans. It has been suggested that glycosylation is important for influenza A virus adaptation in humans [42] and gaining a potential glycosylation site on the HA head at residue 162 may have given clade 6B.1 viruses an evolutionary advantage. Clade 6B.1 viruses also encode an NS1-E125D mutation which increases NS1 interactions with cellular cleavage and polyadenylation factor 30 (CPSF30) [43]. The NS1-E125D mutation of 6B.1 viruses has been suggested to be an important marker for virus adaption to humans [7].

Our study is limited in several ways. The different race distributions between our surveillance sites make it difficult to adjust for in this and any study conducted across different geographic regions. The identification of 4 genetic subgroups reduced the power of our study to detect demographic and clinical differences in our populations. Increasing surveillance and whole genome sequencing efforts could dramatically improve the ability to identify novel virus variants that are causing altered disease in humans. Given these limitations, it is still clear that there is significant H1N1 genetic diversity across the two surveillance sites and this genetic diversity contributes to the differing number of influenza cases and disease severity. Influenza is a global disease, but surveillance at the local level is needed to fully understand virus circulation and disease potential.

## Conflicts of interests

The authors declare no conflict of interest.

## Author contributions

H.L., R.R., K.F.C., and A.P. conceived of the experimental questions; Y.N.G. performed phylogenetic tree analyses; K.S.S., and K.F.C. ran demographic and clinical analyses; T.M., J.E., and P.T. sequenced samples and cleaned the raw data; Z.Y.L., M.L., and L.S. oversaw enrollment and collection of samples from patients; H.L., Y.N.G., K.S.S., and K.F.C. analyzed data; H.L., and A.P. wrote the manuscript; all authors edited and reviewed the manuscript prior to submission.

## Funding

This work was supported by the NIH/NIAID Center of Excellence in Influenza Research and Surveillance contract HHS N2772201400007C (R.R., K.F.C., and A.P.), the Richard Eliasberg Family Foundation and the Research Center for Emerging Viral Infections from The Featured Areas Research Center Program within the framework of the Higher Education Sprout Project by the Ministry of Education (MOE) in Taiwan and the Ministry of Science and Technology (MOST), Taiwan MOST 109-2634-F-182-001 (Y.N.G.).

## Acknowledgements

The authors thank the patients who enrolled and participated in the JH-CEIRS surveillance study. We are grateful for the efforts of the clinical coordination team at JHMI who collected samples. We thank members of the Pekosz lab for feedback on this work.

---

**Supplemental Figure 1.**
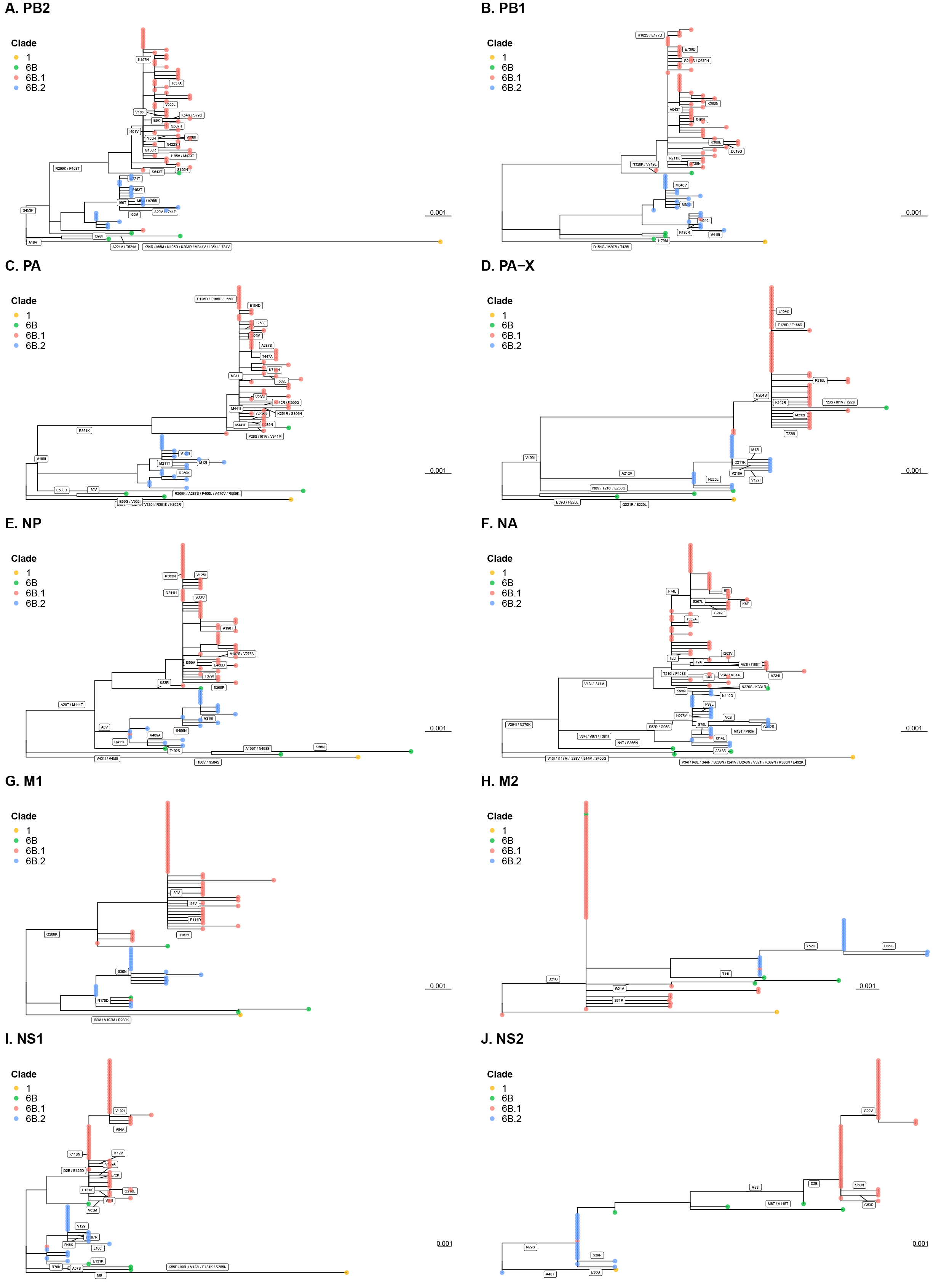
All other segments than HA of H1N1pdm viruses in the 2015-16 season were differentially evolving toward either the clade 6B.1 or 6B.2. Phylogenetic trees of the PB2, PB1, PA, PA-X, NP, NA, M1, M2, NS1 and NS2 coding sequences (subfigures from A to J, respectively) of 73 H1N1pdm viruses of the surveillance were generated using RAxML under GTRGAMMA model with 1000 bootstrap replicates and visualized using the ggtree R package. Branch tips were colored by HA clades, and amino acid mutations on branch were shown. Reference sequences of H1N1pdm vaccine strains, A/California/07/2009 (clade 1), and A/Michigan/45/2015 (clade 6B.1) were included. NP coding sequence numbering begins from upstream start codon.

**Supplemental Table 1.** GenBank accession numbers of each segment of 73 H1N1pdm strains in this study.

| Strain name | HA clade | PB2 | PB1 | PA | HA | NP | NA | M | NS |
| --- | --- | --- | --- | --- | --- | --- | --- | --- | --- |
| A/Baltimore/0008/2016 | 6B.1 | KY487410 | KY487672 | KY487343 | KY487698 | KY487491 | KY487604 | KY487576 | KY487127 |
| A/Baltimore/0026/2016 | 6B.1 | KY949909 | KY949764 | KY950028 | KY950126 | KY949939 | KY949753 | KY950197 | KY949800 |
| A/Baltimore/0065/2016 | 6B.1 | KY950125 | KY949852 | KY950143 | KY950052 | KY949710 | KY949880 | KY950094 | KY950041 |
| A/Baltimore/0076/2016 | 6B.1 | KY487315 | KY487375 | KY487419 | KY487689 | KY487608 | KY487361 | KY487616 | KY487333 |
| A/Baltimore/0077/2016 | 6B.1 | KY949732 | KY949997 | KY949930 | KY950108 | KY949912 | KY949844 | KY950209 | KY950085 |
| A/Baltimore/0088/2016 | 6B | KY487517 | KY487300 | KY615534 | KY487140 | KY487313 | KY487269 | KY487124 | KY487551 |
| A/Baltimore/0091/2016 | 6B.1 | KY487652 | KY487289 | KY487640 | KY615553 | KY615415 | KY487668 | KY487464 | KY487393 |
| A/Baltimore/0092/2016 | 6B.1 | KY487138 | KY487194 | KY615404 | KY487543 | KY487486 | KY487463 | KY487409 | KY487653 |
| A/Baltimore/0096/2016 | 6B.1 | KY487098 | KY615465 | KY487452 | KY615388 | KY487164 | KY487401 | KY487428 | KY487191 |
| A/Baltimore/0101/2016 | 6B.1 | KY487518 | KY487065 | KY487522 | KY487503 | KY487259 | KY615395 | KY487263 | KY487762 |
| A/Baltimore/0103/2016 | 6B.1 | KY487208 | KY487397 | KY487290 | KY487506 | KY487341 | KY487256 | KY487384 | KY487201 |
| A/Baltimore/0104/2016 | 6B.1 | KY487282 | KY487502 | KY487344 | KY487646 | KY487471 | KY487475 | KY487605 | KY487212 |
| A/Baltimore/0107/2016 | 6B.1 | KY487556 | KY487562 | KY487722 | KY487472 | KY487183 | KY487084 | KY487451 | KY487439 |
| A/Baltimore/0110/2016 | 6B.1 | KY487731 | KY615501 | KY487168 | KY615408 | KY487185 | KY615483 | KY487645 | KY487752 |
| A/Baltimore/0118/2016 | 6B.1 | KY615383 | KY615428 | KY615505 | KY615604 | KY615376 | KY615516 | KY487552 | KY615550 |
| A/Baltimore/0122/2016 | 6B.1 | KY615526 | KY615507 | KY487237 | KY615559 | KY615489 | KY615491 | KY487461 | KY615412 |
| A/Baltimore/0124/2016 | 6B.1 | KY487667 | KY487566 | KY487101 | KY487575 | KY487158 | KY487434 | KY615563 | KY487666 |
| A/Baltimore/0125/2016 | 6B.1 | KY487348 | KY487415 | KY487427 | KY487453 | KY487458 | KY487175 | KY487215 | KY487497 |
| A/Baltimore/0158/2016 | 6B.1 | KY487156 | KY615599 | KY487293 | KY487430 | KY615592 | KY487539 | KY487525 | KY487281 |
| A/Baltimore/0163/2016 | 6B.1 | KY487433 | KY487770 | KY487188 | KY487592 | KY487572 | KY487523 | KY615557 | KY487524 |
| A/EastBaltimore/0103/2016 | 6B | KY487534 | KY487611 | KY487704 | KY487656 | KY487615 | KY487392 | KY487465 | KY487599 |
| A/EastBaltimore/0107/2016 | 6B.1 | KY487621 | KY487721 | KY487292 | KY487123 | KY487223 | KY487136 | KY487240 | KY487116 |
| A/EastBaltimore/0119/2016 | 6B.1 | KY487252 | KY487758 | KY487325 | KY615447 | KY487381 | KY487391 | KY487596 | KY487498 |
| A/EastBaltimore/0141/2016 | 6B.1 | KY615562 | KY615541 | KY615512 | KY487527 | KY487701 | KY487628 | KY487529 | KY487696 |
| A/Linkou/0029/2015 | 6B.2 | KY949707 | KY950139 | KY949697 | KY950184 | KY950083 | KY949706 | KY950178 | KY950073 |
| A/Linkou/0030/2015 | 6B | KY949913 | KY950117 | KY950138 | KY949802 | KY949767 | KY949760 | KY949954 | KY949636 |
| A/Linkou/0032/2015 | 6B.1 | KY949925 | KY949664 | KY950068 | KY949823 | KY950112 | KY949903 | KY949646 | KY950070 |
| A/Linkou/0039/2015 | 6B.1 | KY949951 | KY949669 | KY949719 | KY949680 | KY949722 | KY949743 | KY949988 | KY950158 |
| A/Linkou/0046/2016 | 6B.1 | KY949658 | KY950088 | KY949714 | KY949679 | KY949884 | KY949888 | KY949993 | KY950131 |
| A/Linkou/0047/2016 | 6B.2 | KY949962 | KY950065 | KY949782 | KY949975 | KY950049 | KY949999 | KY949696 | KY949731 |
| A/Linkou/0049/2016 | 6B.1 | KY950066 | KY950082 | KY950071 | KY950223 | KY949703 | KY949738 | KY949901 | KY950106 |
| A/Linkou/0050/2016 | 6B.2 | KY949917 | KY949704 | KY949660 | KY949736 | KY950061 | KY950170 | KY949787 | KY950194 |
| A/Linkou/0057/2016 | 6B.1 | KY487199 | KY615531 | KY487496 | KY615390 | KY615380 | KY487310 | KY487581 | KY487584 |
| A/Linkou/0065/2016 | 6B.1 | KY487225 | KY487445 | KY487071 | KY487702 | KY487404 | KY487488 | KY487321 | KY487373 |
| A/Linkou/0068/2016 | 6B.1 | KY487554 | KY487719 | KY487387 | KY615570 | KY487505 | KY615429 | KY487610 | KY487555 |
| A/Linkou/0077/2016 | 6B.2 | KY487278 | KY615572 | KY487598 | KY487687 | KY487372 | KY487383 | KY487283 | KY487087 |
| A/Linkou/0084/2016 | 6B.2 | KY487285 | KY615373 | KY487481 | KY487767 | KY487487 | KY487099 | KY487376 | KY487500 |
| A/Linkou/0093/2016 | 6B.1 | KY487658 | KY487319 | KY487260 | KY487613 | KY487717 | KY487623 | KY487380 | KY487067 |
| A/Linkou/0097/2016 | 6B.1 | KY487417 | KY615513 | KY615499 | KY615425 | KY615535 | KY487639 | KY487328 | KY487213 |
| A/Linkou/0098/2016 | 6B.1 | KY487303 | KY615539 | KY487195 | KY615441 | KY487327 | KY487137 | KY487294 | KY487720 |
| A/Linkou/0099/2016 | 6B.1 | KY487476 | KY487764 | KY487131 | KY487353 | KY487597 | KY487197 | KY487068 | KY487072 |
| A/Taipei/0008/2015 | 6B.2 | KY487634 | KY487510 | KY487631 | KY487478 | KY487715 | KY487143 | KY487364 | KY487755 |
| A/Taipei/0015/2015 | 6B.1 | KY487198 | KY487557 | KY487423 | KY487273 | KY615585 | KY487485 | KY487507 | KY487200 |
| A/Taipei/0018/2016 | 6B.1 | KY487754 | KY487607 | KY487690 | KY487725 | KY487167 | KY487501 | KY487692 | KY487378 |
| A/Taipei/0021/2016 | 6B.1 | KY487085 | KY487345 | KY487106 | KY487147 | KY615436 | KY487459 | KY487339 | KY487211 |
| A/Taipei/0025/2016 | 6B.2 | KY487148 | KY487326 | KY487145 | KY487209 | KY487275 | KY487484 | KY487119 | KY487362 |
| A/Taipei/0030/2016 | 6B.1 | KY615580 | KY487726 | KY487079 | KY487336 | KY487277 | KY487553 | KY487489 | KY487386 |
| A/Taipei/0032/2016 | 6B.2 | KY487304 | KY487371 | KY487083 | KY487330 | KY487239 | KY487718 | KY487511 | KY487663 |
| A/Taipei/0037/2016 | 6B.2 | KY487264 | KY487441 | KY487738 | KY487649 | KY487232 | KY615564 | KY487243 | KY487349 |
| A/Taipei/0040/2016 | 6B.1 | KY487411 | KY487538 | KY615496 | KY487744 | KY487241 | KY487229 | KY487089 | KY487257 |
| A/Taipei/0045/2016 | 6B | KY487296 | KY487456 | KY487367 | KY487602 | KY487178 | KY487714 | KY487426 | KY487565 |
| A/Taipei/0046/2016 | 6B.1 | KY487740 | KY487531 | KY487560 | KY487436 | KY487504 | KY487170 | KY487734 | KY487635 |
| A/Taipei/0050/2016 | 6B.2 | KY487526 | KY487474 | KY487110 | KY487591 | KY487312 | KY487102 | KY487746 | KY487111 |
| A/Taipei/0054/2016 | 6B.1 | KY487736 | KY487169 | KY487299 | KY487403 | KY487184 | KY487247 | KY487449 | KY487258 |
| A/Taipei/0055/2016 | 6B.2 | KY487595 | KY487249 | KY487218 | KY487233 | KY487660 | KY487733 | KY487712 | KY487528 |
| A/Taipei/0059/2016 | 6B.2 | KY487320 | KY487118 | KY615435 | KY615586 | KY615461 | KY487324 | KY487648 | KY487291 |
| A/Taipei/0061/2016 | 6B.1 | KY487379 | KY487546 | KY487732 | KY487248 | KY487224 | KY487350 | KY487636 | KY487360 |
| A/Taipei/0063/2016 | 6B.2 | KY487629 | KY487090 | KY615394 | KY487365 | KY487571 | KY487230 | KY487516 | KY487174 |
| A/Taipei/0067/2016 | 6B.2 | KY487311 | KY615407 | KY487470 | KY487709 | KY487388 | KY487675 | KY487255 | KY487226 |
| A/Keelung/0011/2015 | 6B.1 | KY487600 | KY487113 | KY487166 | KY487251 | KY487574 | KY487437 | KY487699 | KY487593 |
| A/Keelung/0013/2015 | 6B.1 | KY950069 | KY949665 | KY949995 | KY949643 | KY950031 | KY950092 | KY949803 | KY950115 |
| A/Keelung/0027/2016 | 6B.2 | KY487512 | KY487713 | KY487086 | KY487340 | KY487676 | KY487231 | KY487134 | KY487418 |
| A/Keelung/0029/2016 | 6B.1 | KY615478 | KY487334 | KY615481 | KY487499 | KY487769 | KY615400 | KY487400 | KY487314 |
| A/Keelung/0031/2016 | 6B.2 | KY487363 | KY487573 | KY615579 | KY615480 | KY487589 | KY487204 | KY487276 | KY487444 |
| A/Keelung/0038/2016 | 6B.2 | KY487483 | KY487739 | KY487614 | KY487655 | KY487268 | KY487466 | KY487091 | KY487129 |
| A/Keelung/0039/2016 | 6B.1 | KY949952 | KY949976 | KY950181 | KY949887 | KY949948 | KY949872 | KY949898 | KY949946 |
| A/Keelung/0042/2016 | 6B.2 | KY487244 | KY487186 | KY487187 | KY487695 | KY487274 | KY615567 | KY487399 | KY615457 |
| A/Keelung/0046/2016 | 6B.1 | KY487665 | KY487761 | KY615468 | KY487149 | KY487406 | KY487271 | KY487245 | KY487217 |
| A/Keelung/0053/2016 | 6B.1 | KY487594 | KY487609 | KY487684 | KY487727 | KY487088 | KY487181 | KY487177 | KY487748 |
| A/Keelung/0055/2016 | 6B.1 | KY949741 | KY949863 | KY950220 | KY950148 | KY949733 | KY949761 | KY949653 | KY950042 |
| A/Keelung/0056/2016 | 6B.1 | KY487082 | KY487495 | KY615527 | KY487617 | KY487368 | KY487416 | KY487443 | KY487165 |
| A/Keelung/0065/2016 | 6B.2 | KY487674 | KY487216 | KY615378 | KY487219 | KY487567 | KY615459 | KY487075 | KY487579 |
| A/Keelung/0068/2016 | 6B.1 | KY487114 | KY487742 | KY487180 | KY615443 | KY487374 | KY487728 | KY487105 | KY487228 |

**Supplemental Table 2.** Amino acid differences (H1 numbering) of HA gene in 6B.1 and 6B.2 viruses of Baltimore and northern Taiwan in the 2015-16 season

| **Vaccine/ Surveillance strains\*** | **HA1-84** | **HA1-152** | **HA1-162** | **HA1-173** | **HA1-209** | **HA1-216** | **HA1-223** | **HA1-261** | **HA2-164** | **HA2-174** |
| --- | --- | --- | --- | --- | --- | --- | --- | --- | --- | --- |
| A/California/7/2009 | S | V | S | V | T<sup>&</sup> | I | R<sup>&</sup> | A | E | D |
| A/Michigan/45/2015 | N | V | N<sup>Gly+</sup> | V | M<sup>&</sup> | T | R<sup>&</sup> | A | E | D |
| 6B.1 parental (n=26) | N | V | N<sup>Gly+</sup> | V | K | T | Q | A | E | D |
| 6B.1 variant (n=23) | N | V | N<sup>Gly+</sup> | V | K | T | Q | A | E | D |
| 6B.2 variant-1 (n=6) | S | T | S | I | K | I | Q | S | G | E |
| 6B.2 variant-2 (n=13) | S | T | S | I | K | I | Q | A | G | E |
| 6B.1 reassortant (n=1) | N | V | N<sup>Gly+</sup> | V | K | T | Q | A | E | D |

\* Twenty-six strains of 6B.1 parental group include 22 Baltimore and 4 northern Taiwan strains. 6B.1 variant, 6B.2 variant-1 and -2, and 6B.1 reassortant strains are all isolated from northern Taiwan.
<sup>Gly+</sup> Amino acid changes lead to gain of potential N-linked glycosylation site.
<sup>&</sup> Mutations of clade 1 A/California/7/2009 vaccine X-179A or clade 6B.1 A/Michigan/45/2015 vaccine X275. The HA1 sequences of clinical isolates of A/California/7/2009 and A/Michigan/45/2015 were 209K and 223Q.

**Supplemental Table 3.** Amino acid differences of NA gene in 6B.1 and 6B.2 viruses of Baltimore and northern Taiwan in the 2015-16 season

| **Vaccine/ Surveillance strains\*** | **NA-13** | **NA-34** | **NA-52** | **NA-67** | **NA-74** | **NA-96** | **NA-314** | **NA-381** |
| --- | --- | --- | --- | --- | --- | --- | --- | --- |
| A/California/7/2009 | V | I | S | V | F | G | I | T |
| A/Michigan/45/2015 | I | V | S | V | F | G | M | T |
| 6B.1 parental (n=26) | I | V | S | V | F | G | M | T |
| 6B.1 variant (n=23) | I | V | S | V | L | G | M | T |
| 6B.2 variant-1 (n=6) | V | I | S | I | F | G | I | I |
| 6B.2 variant-2 (n=13) | V | I | R<sup>Gly-</sup> | I | F | S | I | I |
| 6B.1 reassortant (n=1) | V | I | R<sup>Gly-</sup> | I | F | S | I | I |

\* Twenty-six strains of 6B.1 parental group include 22 Baltimore and 4 northern Taiwan strains. 6B.1 variant, 6B.2 variant-1 and -2, and 6B.1 reassortant strains are all isolated from northern Taiwan.
<sup>Gly-</sup> Amino acid changes lead to loss of potential N-linked glycosylation site.

**Supplemental Table 4.** Amino acid differences of internal genes in 6B.1 and 6B.2 viruses of Baltimore and northern Taiwan in the 2015-16 season

| **Vaccine/ Surveillance strains\*** | **PB2-**<br>**299** | **PB2-**<br>**453** | **PA-**<br>**361** | **PA-X-**<br>**212** | **NP-**<br>**6<sup>&</sup>** | **M1-**<br>**30** | **M1-**<br>**208** | **M2-**<br>**52** | **NS1-**<br>**2** | **NS1-**<br>**125** | **NS2-**<br>**2** | **NS2-**<br>**22** | **NS2-**<br>**83** |
| --- | --- | --- | --- | --- | --- | --- | --- | --- | --- | --- | --- | --- | --- |
| A/California/7/2009 | R | S | K | A | A | S | Q | Y | D | E | D | G | M |
| A/Michigan/45/2015 | R | P | K | A | A | S | K | Y | E | D | E | G | I |
| 6B.1 parental (n=26) | K | T | K | A | A | S | K | Y | E | D | E | G | I |
| 6B.1 variant (n=23) | K | T | K | A | A | S | K | Y | E | D | E | V | I |
| 6B.2 variant-1 (n=6) | R | P | R | V | V | S | Q | Y | D | E | D | G | M |
| 6B.2 variant-2 (n=13) | R | P | R | V | V | N | Q | C | D | E | D | G | M |
| 6B.1 reassortant (n=1) | K | T | K | A | V | N | Q | C | D | E | D | G | M |

\* Twenty-six strains of 6B.1 parental group include 22 Baltimore and 4 northern Taiwan strains. 6B.1 variant, 6B.2 variant-1 and -2, and 6B.1 reassortant strains are all isolated from northern Taiwan.
<sup>&</sup> NP coding sequence numbering begins from upstream start codon.

## Notes

### Competing Interest Statement

The authors have declared no competing interest.

### Summary of Updates

New title, updated phylogenetic trees and tables of amino acid changes.

